# Collective motion as a distinct behavioural state of the individual

**DOI:** 10.1101/2020.06.15.152454

**Authors:** Daniel Knebel, Ciona Sha-ked, Noa Agmon, Gil Ariel, Amir Ayali

**Affiliations:** School of Zoology, Faculty of Life Sciences, Tel Aviv University, Tel Aviv, 6997801, Israel; Department of Computer Science, Bar-Ilan University, Ramat-Gan, 5290002, Israel; Department of Mathematics, Bar Ilan University, Ramat-Gan, 5290002, Israel; Sagol School of Neuroscience, Tel Aviv University, Tel Aviv, 6997801, Israel

**Author notes:** Lead Contact author Amir Ayali.

## Abstract

The collective motion of swarms depends on adaptations at the individual level. We explored these and their effects on swarm formation and maintenance in locusts. The walking kinematics of individual insects were monitored under laboratory settings, before, as well as during collective motion in a group, and again after separation from the group. It was found that taking part in collective motion induced in the individual unique behavioural kinematics, suggesting the existence of a distinct behavioural mode that we term a “collective-motion-state”. This state, characterized by behavioural adaptation to the social context, is long lasting, not induced by crowding per-se, but only by experiencing collective motion. Utilizing computational models, we show that this adaptability increases the robustness of the swarm. Overall, our findings suggest that collective-motion is not only an emergent property of the group, but also depends on a behavioural mode, rooted in endogenous mechanisms of the individual.

## Introduction

The ability to form groups that move collectively is a key behavioural feature of many species (Sumpter, 2006; Ward and Webster, 2016), assumed to increase the survival of both individuals and groups (Be’er and Ariel, 2019; Yang and Schmickl, 2019). Collectively moving organisms, however, differ in the levels of peer-to-peer interactions, ranging from minimal cooperation to complex social behaviours (Attanasi et al., 2014; Cavagna et al., 2010). Furthermore, endogenous differences among individuals, heterogenic environments, and variability in the interactions between the individual and its direct environment, are all sources of variance that may affect the coordinated behaviour of the collective. Accordingly, it is not clear how synchronized collective motion constitutes such a robust phenomenon, maintaining its form across various group sizes and densities, and under heterogeneous and unpredictable environmental conditions.

One of the most interesting, albeit disastrous, examples of collective motion is that of the marching of locusts. These insects swarm in groups of millions, migrating in mass across large distances, devastating vegetation and agriculture (Ayali, 2019; Cullen et al., 2017; Zhang et al., 2019). In the context of social interactions, locust swarming is characterized by a minimal level of cooperation between individuals: collectivity, which is based on local interactions, is mostly manifested in alignment among neighbouring individuals and in maintaining the overall movement in the same general direction (e.g. Ariel et al., 2014a; Bazazi et al., 2008). Nonetheless, the locust swarming phenomenon is extremely robust, with huge swarms demonstrating moderate to high collectivity on huge scales (up to 6-7 orders of magnitudes), in terms of both the number of animals and their spatio-temporal distribution (Ellis and Ashall, 1957; Uvarov, 1977; see further references in Ariel and Ayali, 2015). Thus, locusts exhibit a considerable disparity between little local cooperation and large-scale collectivity.

What is the key to this ability of locust swarms to maintain their integrity? Here, we show by a series of carefully controlled behavioural experiments that collective movement induces an internal switch in the individual gregarious locust, activating a behavioural mode we refer to as a “collective-motion-state”. In this state, the kinematic behaviour of individuals notably differs from that during a non-collective-motion state. It is important to emphasize that both the “collective-motion-state” and the “non-collective-motion-state” are internal states of swarming-gregarious locusts. We are not referring to the well-known solitarious-gregarious phase transition in locusts (Ayali, 2019; Cullen et al., 2017).

How, then, does the collective-motion-state affect the formation and robustness of the swarm? Interestingly, the switch into this state seems to occur rapidly, and in response to coordinated walking. In particular, our experiments indicate that aggregation alone is not sufficient. Switching out of the collective-motion-state occurs over a longer time scale – significantly longer than the typical time scale of normal fluctuations around the swarm typical dynamics. Hence, stochastic fluctuations, typical to swarming behaviour (Algar et al., 2019; Ariel and Ayali, 2015; Escaff et al., 2018) are “smoothed-out”, leading to highly robust dynamics of the swarm collective behaviour, which is in turn beneficial for the swarm integrity.

Using a simplified computer model, we simulated the swarming properties of locust-like agents with different kinematic parameters, representing the different behavioural states. The results support the functional advantages of the collective-motion- state; allowing us to conclude that the collective-motion-state provides an individual-based mechanism that increases the stability of swarms in the presence of fluctuations, preventing the swarm from collapsing.

## Results

The main objective of this report was to examine the behaviour of individual animals upon joining and mostly leaving a group of conspecifics. Here we studied gregarious locusts, reared in dense populations, one developmental stage before becoming adults and developing functional wings (i.e. fifth larval instar). The experiments comprised three consecutive stages, representing different conditions (Fig. 1 and Vid. S1): (1) Isolation stage: a single animal was taken from its highly dense rearing cage, tagged with a barcode, and introduced alone into a ring-shaped arena (outer and inner diameters: 60 cm and 30 cm, respectively); (2) Grouping stage: nine other individually tagged animals were added to the arena; and (3) re-isolation stage: the nine added animals were removed from the arena, leaving the original animal alone. The duration of each stage was one hour, which was enough for the locusts to exhibit their walking kinematics, yet did not cause behavioural changes due to exhaustion, hunger etc. The trajectories of the animals’ were fully reconstructed using a barcode tracking system. The middle 40 minutes of each stage were analysed, as detailed in the Methods section. A range of kinematic statistics was collected in order to classify and compare the locusts’ behaviour in the different stages.

**Figure 1.**
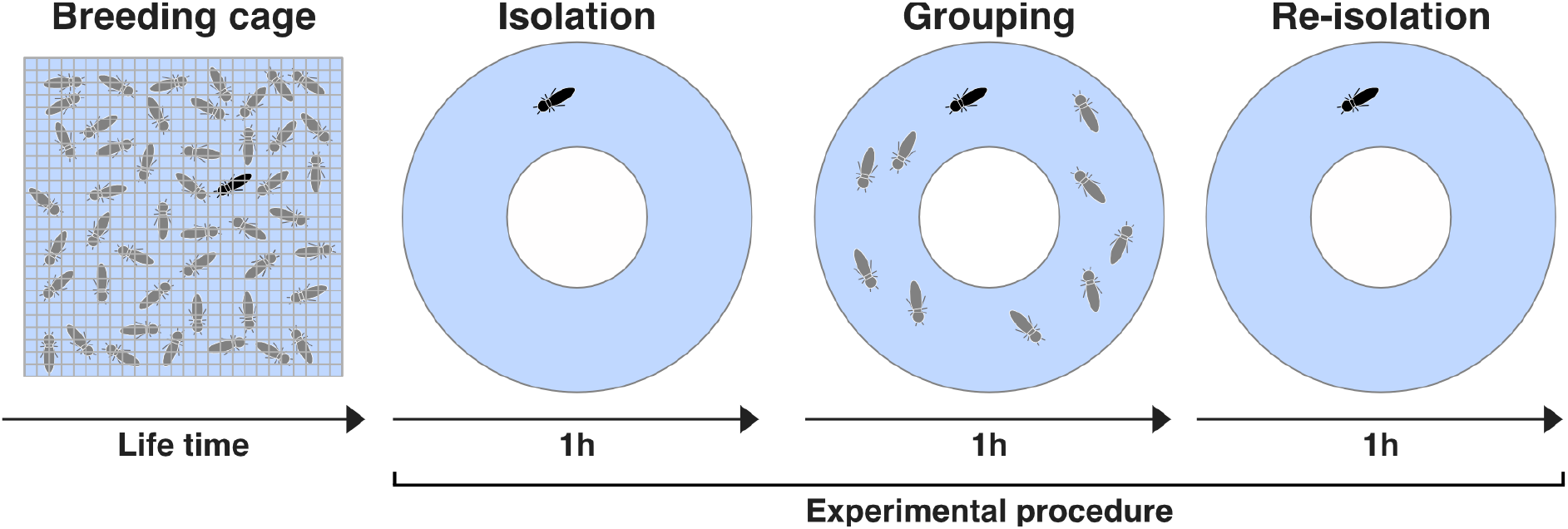
A schematic flow of the experimental procedure. Locusts were reared in high density conditions. The Experiments comprised the following consecutive stages: (1) isolation for one hour in the arena; (2) grouping for one hour; and (3) re-isolation for one hour.

### Swarm formation – validation of collective motion

In order to verify that our grouping conditions were indeed inducing collective motion (swarming), we calculated the synchronization in movement of the grouped animals using the order parameter (see Methods for definition), which is a fundamental estimator for the typical marching behaviour of locusts (e.g. Knebel et al., 2019). The median order parameter in the grouping stage was found to be significantly higher than that obtained for computationally randomized groups (as presented in our previous report, Knebel et al., 2019; medians: 0.632 and 0.239 respectively; Wilcoxon rank-sum test: *p* < 0.001). Consequently, we conclude that the groups in our experimental setup indeed demonstrated swarming and collective motion.

### Kinematic differences among the isolation, grouping, and re-isolation stages

Locusts walk in an intermittent motion pattern (Ariel et al., 2014a; Bazazi et al., 2012), i.e., movement occurs in sequences of alternating walking bouts and pauses. To characterize individual locust kinematics, we measured four parameters: (1) the fraction of time an animal spends walking; (2) the average speed while walking; (3) the average walking bout duration; and (4) the average pause duration. Comparing these values across the three experimental stages, we found several statistically significant differences (Fig. 2). In the following, *p*-values correspond to a Friedman test followed by a multiple comparison test using the Bonferroni method.

**Figure 2.**
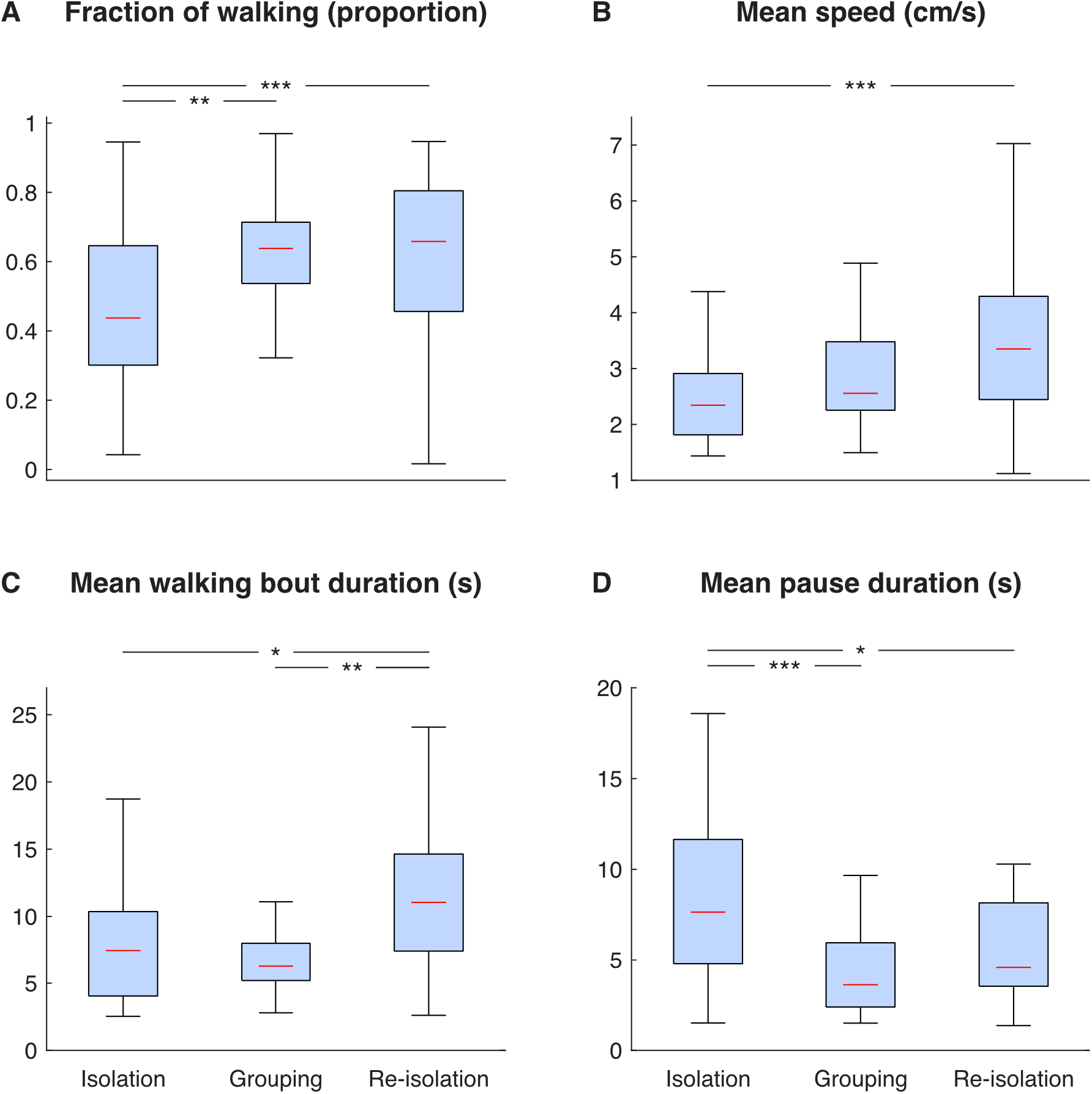
Kinematic changes throughout the three experimental conditions. (a) the fraction of walking, (b) the averaged walking speed, (c) the average duration of walking bouts, and (d) the average duration of pauses, of the traced animals in the isolation, grouping, and re-isolation stages. \**p*<0.05, \*\**p*<0.01, \*\*\**p*<0.001.

We found that when comparing between isolation and grouping, the fraction of time spent walking and the pause durations differed significantly (*p*<0.01 and *p*<0.001, respectively), showing a larger fraction of time walking and shorter pauses while grouping. These findings are in accordance with previous reports (Knebel et al., 2019), and are consistent with the known propensity of locusts to walk more and rest for shorter times while in a swarm (Ariel et al., 2014a; Bazazi et al., 2012, 2008; Knebel et al., 2019). However, our experiments also revealed a new effect of swarming. Comparing the isolation and re-isolation stages, we found that all the studied parameters differed significantly. Specifically, the fraction of time walking, speed, and walking bout duration were all higher in the re-isolation stage, while the pause duration decreased (*p*<0.001, *p*<0.001, *p*<0.05 and *p*<0.05, respectively). Interpreting these parameters together, while also taking into account the low propensity of locusts to turn while walking (or to make a U-turn upon starting to walk; Ariel et al., 2014a), the overall area explored by the locusts was much larger during the second isolation state. Furthermore, comparing the grouping and re-isolation conditions revealed that the walking bout duration increased significantly following re-isolation (*p*<0.01). The data for all other comparison combinations were found not to differ significantly. The exact data points and trends can be found in Fig. S1.

The increase in activity following re-isolation is surprising, and suggests that the marching behaviour of locusts is not dictated by instantaneous or immediate interactions among individuals per-se. Rather, our findings indicate that the interactions with other marching locusts induce a switch to a new internal behavioural state, which outlasts the presence of the swarm. In accordance with our results, we term this internal state the “collective-motion-state”. The different behavioural states are schematically presented in Fig. 3.

**Figure 3.**
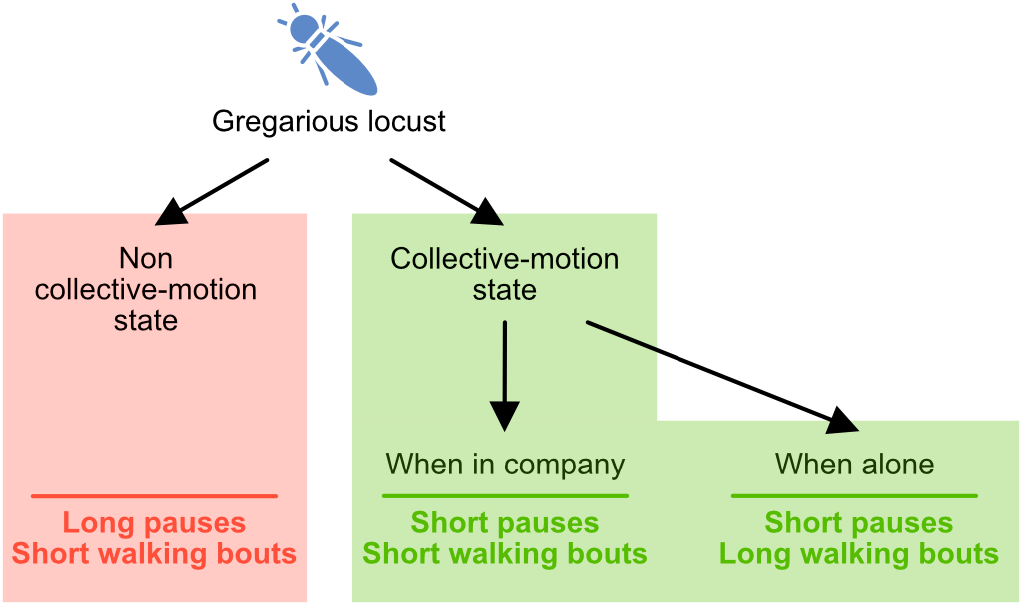
A schematic representation of the behavioural states of the locusts.

In order to verify that the observed behavioural changes indeed represent a transient state, rather than a permanent behavioural modulation, a simple control experiment was performed. In 6 of the experiments, following the re-isolation stage, locusts were returned to their rearing cage (in high crowding conditions without collective motion), and tested the next day again alone in the arena (isolation condition). We found no significant behavioural difference between this latter isolation and the first isolation stage of the previous day. Hence, the collective-motion state is transient. A second series of control experiments (n=6) was performed in order to exclude potential time effects on the locusts’ behaviour due to the duration of the experiments. To this end, locusts were tested in isolation for three consecutive hours. The kinematic analysis procedure described above was then performed separately on three 40 min segments of the 3 h tests and no significant differences were found.

### Consistency of individual behavioural tendencies

Despite the observed major differences in behavioural kinematics among the three experimental stages, we were also interested to know whether there are any correlations between the changing parameters in the three experimental conditions: isolation, grouping and re-isolation. This would indicate that although individuals change their behaviour throughout the experimental stages (Fig. 2), they maintain the relative position in comparison to others, and thus show some consistent individual tendencies. We found that individual behavioural tendencies generally persisted. The fraction of walking, speed, and pause durations all showed high within-individual correlation across the three stages. The walking bout durations, however, were significantly correlated only between the isolation and grouped stages, but not between re-isolation and the other stages. (see Table 1 for numerical details). This suggests that while fraction of walking, speed and pause durations are highly dependent on the animal tested itself, the bout duration during re-isolation cannot be predicted by the previous stages, and is therefore influenced by the social context rather than by the animal’s unique properties.

**Table 1.** Correlation values between kinematic parameters throughout the three experimental conditions. The fraction of walking, the averaged walking speed, the average duration of walking bouts, and the average duration of pauses were tested for correlation across the three experimental conditions: (1) isolation, (2) grouping, and (3) re-isolation. The underlined values mark significant correlations.

|  | *p-Value* | *rho-Value* |
| --- | --- | --- |
| **Fraction of Walking** |  |  |
| Isolation - Grouping | <u>0.000</u> | <u>0.706</u> |
| Isolation - Re-isolation | <u>0.001</u> | <u>0.679</u> |
| Grouping - Re-isolation | <u>0.002</u> | <u>0.638</u> |
| **Speed** |  |  |
| Isolation - Grouping | <u>0.001</u> | <u>0.679</u> |
| Isolation - Re-isolation | <u>0.010</u> | <u>0.561</u> |
| Grouping - Re-isolation | <u>0.002</u> | <u>0.636</u> |
| **Walking bout duration** |  |  |
| Isolation - Grouping | <u>0.002</u> | <u>0.628</u> |
| Isolation - Re-isolation | 0.148 | 0.391 |
| Grouping - Re-isolation | 0.354 | 0.314 |
| **Pause duration** |  |  |
| Isolation - Grouping | <u>0.007</u> | <u>0.580</u> |
| Isolation - Re-isolation | <u>0.003</u> | <u>0.612</u> |
| Grouping - Re-isolation | <u>0.000</u> | <u>0.708</u> |

### Modelling and simulations

The experiments described above demonstrated that individual locusts introduced into a collectively moving swarm undergo a switch into a distinct internal socio-behavioural state. However, whether this change confers a benefit on the swarm formation and maintenance, and if so, of what kind, remained unanswered. To explore this aspect of the collective-motion-state, we developed a model that simulates locust swarms, in which individual kinematics could be manipulated.

To simulate swarms, we used a simplified agent-based model in a square domain with periodic boundaries. Agents were designed as rectangles with a circular receptive field around their centre. The agents’ kinematics were programmed to resemble that of locusts, to move in an intermittent motion (pause-and-go) pattern: at every step of the simulation, each agent made an individual decision whether to walk or stop, based on its current state (walking or stopping) with predefined probabilities (*p*_W_ and *p*_S_, respectively). While moving, the speed was constant. The individual direction of movement was allowed to change only when an agent changes its state from stopping to walking (Ariel et al., 2014a). An agent’s new direction was a weighted sum of its own direction (inertia), the direction of other agents in its visual field, with a short memory (see Rimer and Ariel, 2017 for the importance of memory in pause-and-go simulations), and noise. These were set to generate an order value approximately similar to that obtained in our experiments. See Methods for details and Table S1 for parameter values.

The spatial and temporal scales of the model were set as follows: 1 cm was considered as a distance of 0.1 in the simulated arena, and 1 sec corresponded to 1 simulation step. Thus, we fixed the model dimensions to correspond with real locusts’ size and movement parameters. The size of agents (0.1 × 0.4) maintains the proportions of 5th larval instar locusts. Additionally, the visual range of 1 radius, ten times larger than the agent’s rectangle width, represents the proportional visual field of locusts (range just under 10 cm; Ariel et al., 2014a). The speed was set to 0.25, corresponding to the typical swarming speed shown in Fig 2b. Finally, the size of the arena was set to be 7×7, with 12 agents within. This generated a slightly lower density than in the real experiments, but better mimicked the limited visual field the locusts experienced in our ring shape experimental arena.

We evaluated the effect of the agent’s walking bout and pause durations, controlled by *p*_W_ and *p*_S_, on four statistics that characterize collective motion in the swarm:

1. The order parameter (the size of the average direction vector).
2. Spread (the average distance between all pairs).
3. The number of neighbours (within the field of view, denoted NN).
4. Regrouping time (the average number of steps it takes an agent that has lost all other agents in its receptive field to re-obtain at least one neighbour).

Statistics were averaged over all agents and simulation steps.

Fig. 4 shows the median value over 40 independent repetitions for each parameter set. We found that the order parameter increases with the walking bout duration but decreases with pause durations (Fig. 4B). The spread, on the other hand, is increasing both with walking and pausing durations (Fig. 4C). The NN statistics (Fig. 3D) were in accord with the spread (small spread implies many NN). Its values were similar to that obtained in the real experimental arena (~ 1-1.5), which was slightly denser, as explained above. The regrouping time (Fig. 4E) showed a more complex dependency on the duration of walk and pause durations: it was shortest when both the parameters were low, and longest when the walking bout was low but the pause duration was high. Yet, for mid values of pause duration, high walking bouts durations induced a reduction in the number of steps to regroup. Worth noting is the fact that because agents could only change direction when starting to move (similar to the locusts; Ariel et al., 2014a), the length of walking segments between turns increases with walking durations.

**Figure 4.**
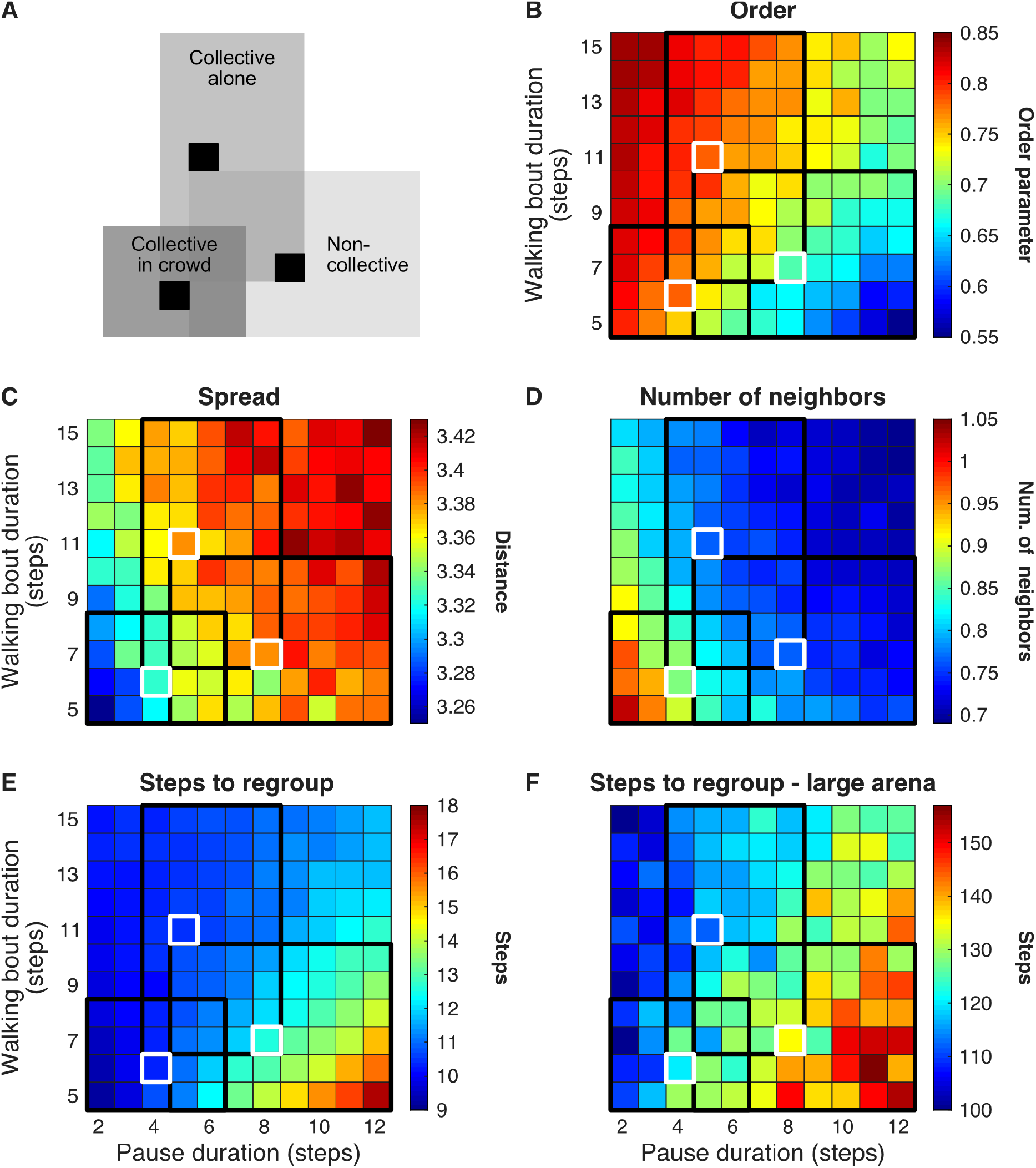
The influence of different walking bout and pause durations on the collectivity parameters in simulated swarms. (A) The areas representing the behavioural states in the next traces, where the black and white frames indicate the inter quartile ranges and medians of each condition, respectively, obtained from Fig. 2C and D. (B) The order parameter, (C) spread, (D) the average number of agents within each agent’s visual field, and (E) the average number of steps to regroup, for different walking bout (rows) and pause (columns) average durations. The arena size is 7×7 with periodic boundaries. (F) The average number of steps to regroup in a large 75×75 arena, with the same number of agents. In (B-E) each coloured box represents the median of 40

In order to relate the simulation and experimental results, the three collective motion states – isolation, grouping and re-isolation – were indicated in Fig. 4 by the tiles corresponding to the inter quartile range of the empirical data presented in Fig. 2C and D. Additionally, we also marked the corresponding median data point. As can be seen, in all of the parameters calculated, the individual behaviour that reflects the collective state in a crowd, improves the coherency and rigidity of the swarm. This important observation suggests a possible benefit for increasing the walking bout duration when a locust is in a collective state, but finds itself alone.

In order to further explore the advantages of a collective motion step in re-isolation scenarios, we increased the size of the arena to 75×75, with the same number of agents, thereby reducing the density considerably, and calculated the steps to regroup parameter (Fig. 3E). We found that for low density, where practically no collective behaviour is present (see Fig. S3), longer walking bouts reduce the time to regroup. Therefore, it is beneficial for an agent that finds itself alone due to a sparse distribution of the swarm to increase its walking bouts duration (even in the cost of decreasing other parameters of collectivity), and thus shorten the time until it reunites with other locusts.

## Discussion

Our findings reported here suggest that, in locusts, the sensory-motor act of collective motion is accompanied by an internal state of the individual locust – a collective-motion-state, that is manifested in specific behavioural kinematics. This state is induced by the experience of synchronous, collective marching. In turn, it has an important role in maintaining the integrity and consistency of the swarm. Below we discuss several key aspects and implications of this finding.

It should be stressed once again that the current study focused on gregarious, crowded-reared locusts only. The described behavioural states should not be confused, therefore, with the well-known and much researched locust density-dependent phase polyphenism (Ayali, 2019; Cullen et al., 2017). Collective motion is limited to the gregarious, swarming and migrating phase. Accordingly, all our experimental animals were taken from our gregarious (crowded-reared) breeding colony, maintained in crowded conditions for many consecutive generations. In their breeding cages, mostly due to the physical constraints and abundance of food, despite experiencing high density, the locusts very rarely, if at all, demonstrate collective motion. Thus, they adopted the collective-motion-state only upon experiencing, and taking part in, collective marching within the experimental arena.

In a recent study (Knebel et al., 2019), we have introduced a comparison between the walking behaviour kinematics of individual gregarious locusts in different social (density) contexts. Our reported findings are reconfirmed and further elucidated here by the results of the initial isolation and grouping stages in our experiments. The novel idea posited here is that these differences represent not only the spatially and temporally immediate social environment and the instantaneous local interactions among locusts. They are also dictated by the effects of an internal state induced by the general experience of collective motion. A fundamental aspect of the concept of the collective-motion-state arises from our findings related to its persistent effect in time: upon re-isolation, the individual locust adopted behavioural kinematics that critically differed from that in the first experimental stage (initial isolation). We also showed that, as expected, the collective-motion-state is transient. If the locust does not experience collective motion for some time, and is then isolated once more, it loses the unique walking-related kinematics it previously adopted in response to the collective motion, i.e. the internal collective-motion-state. The dynamics of this decay were not explored, but are likely to be affected by many external factors, such as the availability of food and the day-night cycle.

The individual locusts in our experiments retained the variability demonstrated in our previous report (Knebel et al., 2019), while demonstrating a second layer of variability or plasticity upon experiencing collective motion, when entering the collective-motion-state. Considerable research has been devoted in recent years to understanding the effect of variability among individuals on the group’s collective behaviour, both experimentally—ranging from bacteria to primates (Benisty et al., 2015; Brown and Irving, 2014; Crall et al., 2016; Dyer et al., 2009; Farine et al., 2017; Fürtbauer and Fry, 2018; Herbert-Read et al., 2013; Jolles et al., 2018; Planas-Sitjà et al., 2015)—and theoretically (Aplin et al., 2014; Ariel et al., 2014b; Calovi et al., 2015; Copenhagen et al., 2016; Guisandez et al., 2017; Jolles et al., 2017; Menzel, 2012; Mishra et al., 2012; see Mar Delgado et al., 2018; Modlmeier et al., 2015; Webster and Ward, 2011 for recent reviews). The interactions between variability in specific aspects of the individuals’ behaviour and group-level processes were found to be complex and, moreover, bidirectional (e.g. Knebel et al., 2019). Variability among individual animals was found to have important consequences for the collective behaviour of the group (e.g. O’shea-Wheller et al., 2017; Szorkovszky et al., 2018).

However, beyond the variability among the individuals composing a group, variability is also expected in the behaviour of the individual animal over time, as it experiences changes in environmental and social conditions. The swarm (or flock, shoal, herd, etc.) is a heterogeneous entity, moving in a heterogeneous environment. The individual is bound on occasion to find itself in different locations within the swarm (e.g. leading edge, at the outskirts, trailing), and it may also find itself separated from the group by natural obstacles (vegetation, rocks and boulders). It is essential for the robustness and consistency of the swarm that throughout these changing conditions, the behaviour of the individual will adapt accordingly, such as to be appropriate for the changing context. For example, if temporarily separated from the core of the swarm, a locust’s walking kinematics should change to support rapid reunion with the group, as reported in both our experimental and simulation findings (e.g. increased fraction of walking and duration of walking bouts). If previously naïve to collective motion, that individual’s kinematics would however be disadvantageous, or even hinder the formation of a swarm.

In Bazazi et al., 2012, the authors suggest that behavioural variability can be explained by the existence of two internal states. Studying single locusts in isolation for eight consecutive hours, they have observed changes in behavioural kinematics that were suggested to result from “internal state behavioural modulation”. The observed variations, however, were merely attributed to changes in “starvation/satiation state”, i.e. as the locust becomes starved, it changes its walking behaviour, searching more vigorously for food. Moreover, they conclude that animals continually switch between the two states on a scale of minutes. The collective-motion-state reported here is, of course, a very different type of internal behavioural state, which is strongly involved with the locust past and current social environment. It may be viewed as a form, or a manifestation of a social carry-over effect (Niemelä and Santostefano, 2015), where a social environment experienced by a focal individual affects aspects of its locomotion behaviour at a later, non-social context. As noted, however, the change in behavioural state described here is induced by collective marching, i.e., a particular mode of social interaction, rather than by aggregation or being around other conspecifics per-se. Moreover, as our simulations show, the enhanced marching displayed in the re-isolation stage is advantageous for maintaining collective swarming - it is still much related to the social context rather than carried-over to a non-social one.

The reported collective-motion-state is also in accord with the overall daily behavioural changes of marching locust swarms. The swarm will spend the night (as well as times of low temperature or other unfavourable climatic conditions) roosting among the vegetation. Upon suitable conditions, after a period of feeding, the locusts will initiate marching – highly synchronized, collective motion. Frequently, when temperature becomes too high around noon, or when dusk arrives, the swarm will again switch to feeding and roosting. These daily patterns call for corresponding changes in the internal behavioural states of the individual locusts and mostly a dedicated collective-motion-state.

In the current work we are cautious in discussing the underlying mechanisms of the behavioural states reported. Although this is beyond the scope of this study, it is clear that these behavioural states represent physiological states. With some confidence, we can speculate about the nature or the physiological mechanisms involved in the demonstrated behavioural states. Behavioural plasticity in locust behaviour has been attributed to various second messengers or neuromodulators, or to the balance among them. Most notable are the biogenic amines (e.g. Serotonin, a prominent bio-amine, was recently reported to inhibit walking behaviour in *Drosophila*; Howard et al., 2019). Hence, it may well be that the (spatial and temporal) immediate social environment affects biogenic amine levels, and these in turn modulate the walking-related behavioural kinematics manifested in the different behavioural states.

Another candidate that may be involved in the collective-motion-state is the locust adipokinetic hormone (AKH). AKH is a metabolic neuropeptide principally known for its mobilization of energy substrates, notably lipid and trehalose, during energy-requiring activities such as flight and locomotion, but also during stress (e.g. Perić-Mataruga et al., 2006). It is well accepted that the metabolic state affects the level of general activity of an organism, and AKHs are reported to stimulate locomotor activity, either directly by way of their activity within the central nervous system (e.g. Wicher, 2007), or via octopamine – a biogenic amine with ample behavioural effects (Verlinden et al., 2010; Yang et al., 2015).

Furthermore, as noted, we have demonstrated here an extended effect of the experience of collective motion. Hence, learning and memory-related mechanisms would also seem to be involved. Again, previous work may suggest some candidate molecules and pathways, including cGMP-dependent protein kinase (PKG), and Protein Kinase A (Geva et al., 2010; Lucas et al., 2010; Ott et al., 2012).

Last, as noted, solitarious phase locusts lack the capacity to demonstrate collective motion, and thus also the collective-motion-state. Accordingly, they differ from gregarious locusts in all the above physiological pathways (bioamines: e.g. Alessi et al., 2014; Cullen et al., 2017; Ma et al., 2015; AKH: Ayali and Pener, 1992; Pener et al., 1997; PKG: Lucas et al., 2010; PKA: Ott et al., 2012). An in-depth investigation of the development of gregarious-like states in solitary locusts should prove to be very enlightening.

A central question is whether a collective (herd, flock, or swarm) is merely a sum of its parts, or a new entity. Most related studies have perceived collectivity as a self-emergent phenomenon, suggesting that new dynamics and behaviour are the result of intricate, multi-body, typically non-linear interactions (e.g. Cucker and Smale, 2007; Vicsek and Zafeiris, 2012). One hidden assumption underlying this perception, is that individuals remain inherently unchanged when isolated or in a crowd. Even studies of heterogeneous swarms, in which conspecifics may differ from each other, still assume consistency in the properties of the individual over time. This is essentially a physical point of view, in the sense that agents/individuals possess certain properties that determine their behaviour across a range of situations. Thus, the collective motion is an emergent property that builds up in particular contexts, such as a sufficiently high local density of animals. This point of view allows, among others, extrapolation from experiments with one, two, or a few animals to large swarms (e.g. Calovi et al., 2015).

Our findings reported here suggest a fundamentally different point of view. We perceive the sensory-motor act of collective motion as accompanied by an internal state – a collective-motion-state that is manifested in specific behavioural kinematics. This state is induced by the experience of synchronous, collective motion. Most importantly, it is *not* induced by spatial aggregation alone. Collectivity, therefore, is not just self-emerging. Rather, the collective-motion-state has an important role in maintaining the integrity and consistency of the swarm. The robustness of the swarm is also a major challenge and requirement in swarming robotics, making the current novel insights applicable and even important also to this emerging field.

In the case of locusts, our far from complete understanding of the swarming phenomenon is also proving crucial for human well-being and survival, as evident from the current devastating locust situation in large parts of Africa and Asia (FAO, 2020). Much scientific attention has been dedicated to the perception, decision-making, and individual kinematics of locusts in a swarm. These efforts have led to various models that attempt to explain the collective behaviour on the basis of local interactions among the individual locusts (see Ariel and Ayali, 2015 for review). The current study is, to the best of our knowledge, the first to include the internal state of the individual locust as an important factor in dictating its behaviour, and in turn affecting the maintenance and the properties of the swarm.

### Limitations of the Study

The study presented here outlines a post-swarming behavioural state of individuals. Clearly, as noted, this state is induced by neuro-chemical changes such as secretion of neuromodulators and/or hormones. Yet, it was beyond of this research to pinpoint the exact neuronal mechanisms involved. Furthermore, the presented model is simplified, and ignores various aspects of locust swarming which might be critical. However, the simplicity is also a virtue of the model, which can be easily generalized to other systems. In addition, although we show that the collective-motion-state is transient, we did not explore its temporal materialization and decline.

## Data and Code Availability

The data will be made available upon request.

## Acknowledgements

This research has been supported by the Israel Science Foundation (research grant 2306/18)

## Author contributions

D.K., G.A., and A.A. designed the study. D.K. and C.S. performed the experiments. D.K. and C.S. analysed the data. D.K. N.A. and G.A constructed the model. D.K. wrote the code, performed simulations and analysed the data. D.K., G.A., and A.A. wrote the manuscript. All authors reviewed and approved the paper.

---

**Supplementary Video S1**:

Experimental procedure (Related to Fig. 1)

