## Supplementary information for "Collective motion as a distinct behavioural state of the individual"

### Supplemental information

#### Figures and tables

| Variable | | Value |
|---|---|---|
| **Simulation parameters** | Num. of agents (N) | **12** |
| | Num. of steps | **1000** |
| | Arena size (periodic boundaries; *L*) | **7 x 7 / 75 x 75** |
| **Agents dimensions** | Size of agents (*a1* x *a2*) | **0.1 x 0.4** |
| | Visual range radius (*r*) | **1** |
| **Movement parameters** | Probability to stop ($p_s$) | **0.5 - 0.917** |
| | Probability to walk ($p_w$) | **0.8 - 0.933** |
| | Speed (*v*) | **0.25** |
| **Direction parameters** | Inertia weight (*q*1) | **0.2** |
| | Current surrounding weight (*q*2) | **0.2** |
| | Memory surrounding weight (*q*3) | **0.3** |
| | Memory length (of surrounding memory) | **5** |
| | Random vector weight (*q*4) | **0.3** |

**Table S1. Model parameters.**

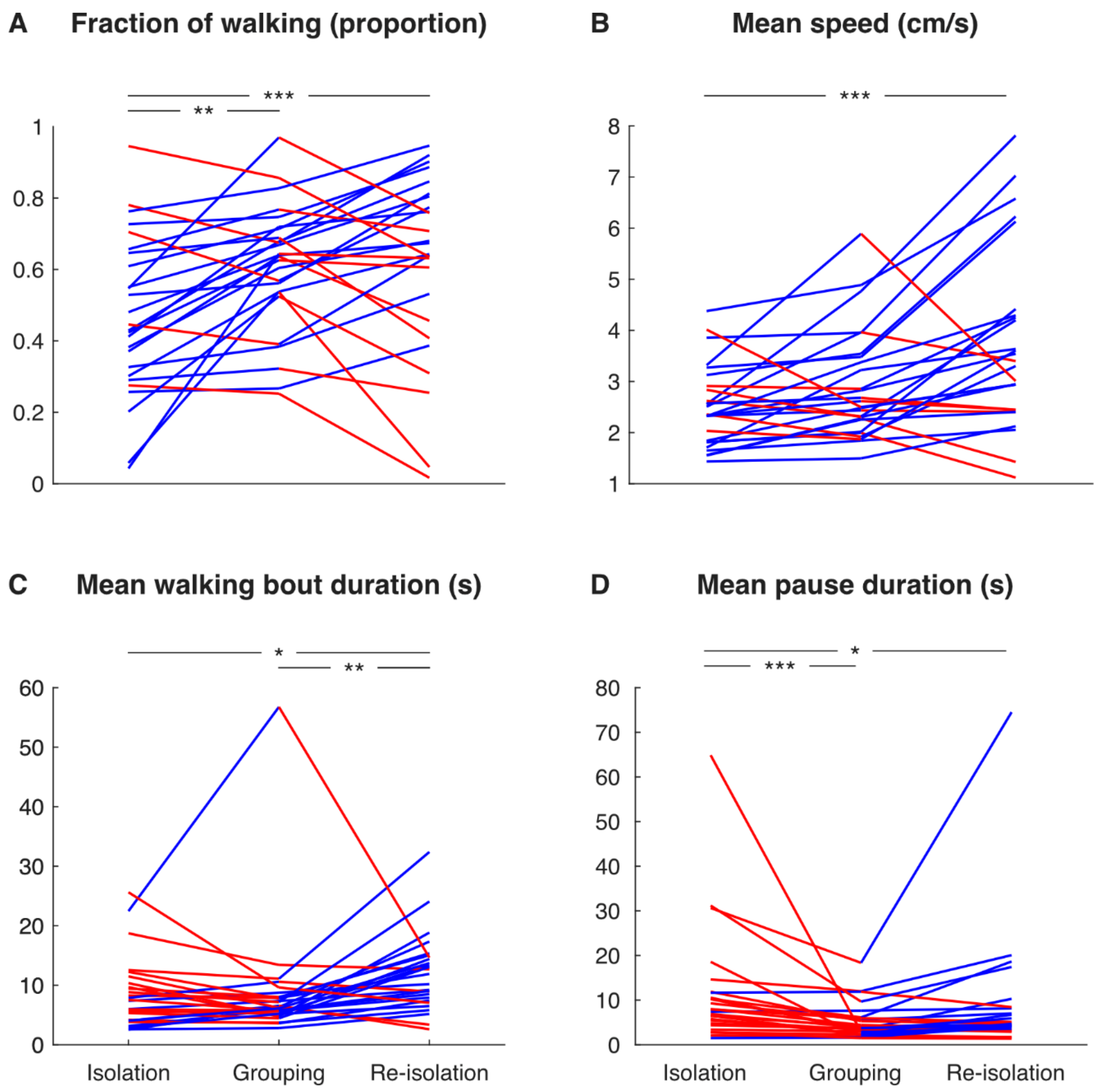


**Fig. S1. Kinematic changes throughout the three experimental conditions – exact data points and trends.** (A) the fraction of walking, (B) the averaged walking speed, (C) the average duration of walking bouts, and (D) the average duration of pauses, of the traced animals in the isolation, grouping, and re-isolation stages. Blue lines represent increase and red lines decrease between consequent experimental conditions of individual animals (n=26). *$p$<0.05, **$p$<0.01, ***$p$<0.001.

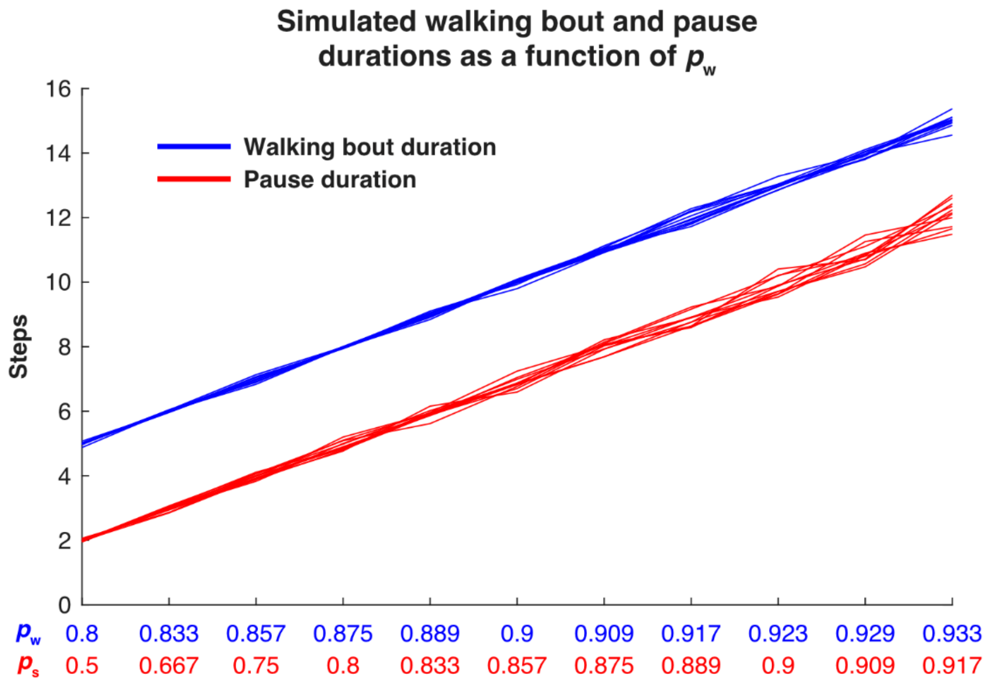


**Figure S2. Walking and pause durations for different $p_W$ values.** The median bout and pause durations presented in Fig. 4, arranged by their corresponding $p_W$ and $p_S$ (blue and red, respectively).

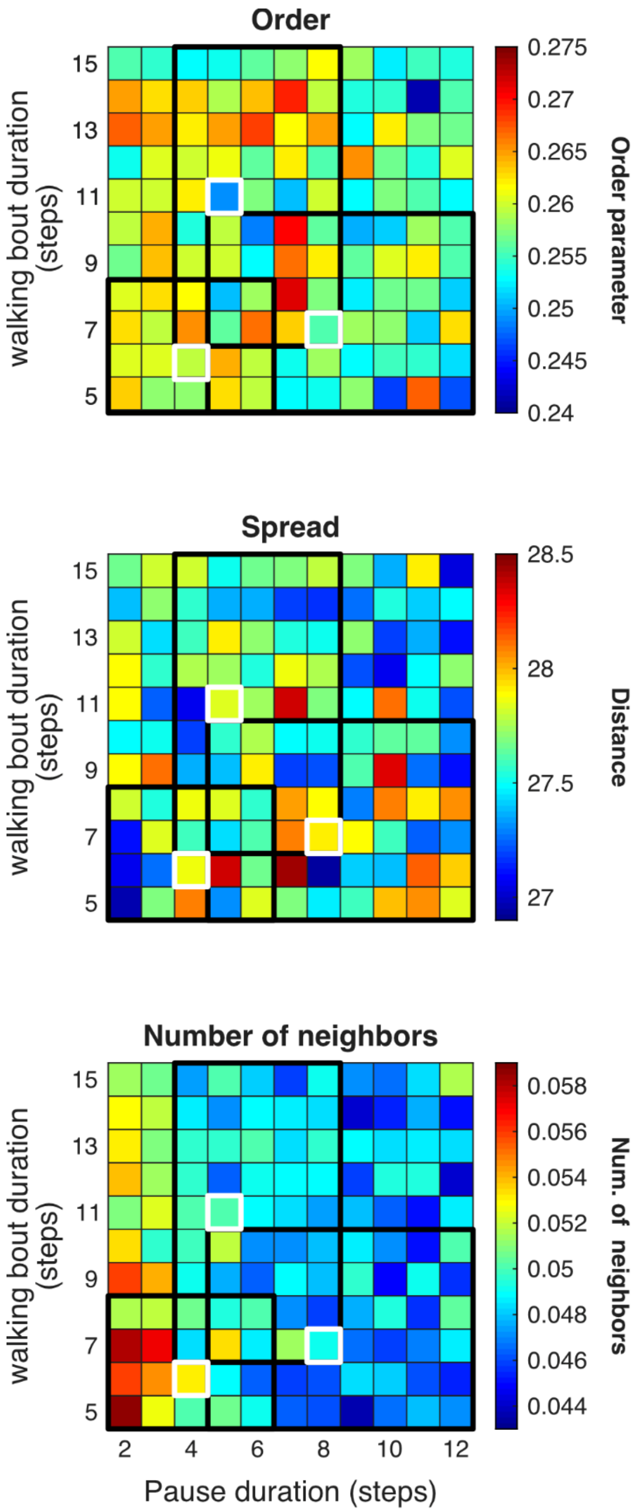


**Figure S3. The influence of different walking bouts and pauses duration on the collectivity parameters of simulated swarms in an arena of 75x75.** The order parameter, spread and the average number of agents within each agent's visual field for different walking bouts (rows) and pauses (columns) durations. Notice that the low numbers and minor changes between the boxes indicate that there was minimal to no collective behaviour in the 75x75 sized arena. Each box represents the median of 100 simulations.

#### Transparent Methods

##### Animals

The empirical experiments were conducted on gregarious desert locusts, *Schistocerca gregaria* (Forsskål), bred at Tel Aviv University, School of Zoology. The animals were kept at high density: over 100 animals in cages of 60 l, under a 12 h:12 h light/dark regime, a temperature of 30℃, and 30-65 % humidity. Animals were fed daily with wheat seedlings and dry oats. All experimental locusts were the offspring of many generations of gregarious locusts reared in these conditions.

##### Experimental setup

A ring-shaped arena was used for the assessment of the behaviours of individuals and swarms. It comprised blue plastic walls (60 cm diameter and 55 cm high) with an inner central cone dome (30 cm diameter). The bottom of the structure was covered with a thin layer of Fluon (Whitford Plastics Ltd., Runcorn, UK), in order to prevent locusts from climbing. A white paper covered the floor of the arena and was changed before each experiment. Light sources around the arena lit its floor evenly. A video camera (Sony FDR-AXP35: 4K Ultra HD) was positioned above the arena and filmed the locusts throughout the experiment.

##### Monitoring the locust movements

Prior to the experiments, each of the experimented locusts was tagged with a unique barcode glued to the dorsal side of its prothorax using a drop of Epoxy glue. Using the video footage at 25/3 frames per second, the trajectory of each animal was reconstructed offline using BugTag software (Robiotec Ltd., Israel). To this we added a custom-designed multiple-target tracking and a trajectory-smoothing method (for details: Ariel et al., 2014a). briefly, segments in which tags were not identified (less than 5 cm or 25 s) by the system positions were interpolated. We thus obtained the positions of the tags' centre of mass relative to the arena centre, at a final resolution of 2-3 pixels (ca. 0.5 mm), for about 99% of the video frames. In order to reach 100% recognition, we completed the analysis manually for the remaining frames. In three out of the 26 experiment the tracking system failed to recognise 1-2 animals in the grouping stage. Therefore, the order parameter was calculated based on the recognized animals only. These miss-identifications, which only slightly affect the order parameter calculated in the grouping stage, have no influence on any of the kinematic statistics in any of the stages (that only describe the focal animal).

**Experimental procedure**

In the morning of each experiment day, 10 locusts were taken from their cage and individually tagged. Thereafter, one locust was randomly chosen and placed in the arena alone for one hour. The other 9 locusts were then added for an additional hour; after which they were removed. The remaining locust was then filmed for another hour.

**Analysed parameters**

Following the recognition process, we analysed the middle 40 min of each of the 3 hours of the experiment using MATLAB (MathWorks, Natick, MA, USA). The following parameters were calculated:

1. Order parameter – the absolute value of the average direction of movement of the walking animals (clockwise denoted as -1 and anticlockwise as +1), averaged for all the frames.

2. Fraction of walking – the number of frames an animal walked divided by the total number of analysed frames.
3. Detection of walking bouts – a walking period was defined as walking at a speed above 0.25 cm/sec for more than one third of a second. This double threshold allowed us to overcome the limitation of the tracking system and avoid recognizing very small movements as real walking.
4. Speed – averaged for all experimental animals and analysed frames when the animal walked and the speed differed from zero.
5. Walking bout duration – the time an animal moved continuously between one pause to the next, averaged for all walking bouts.
6. Pause duration – the time an animal did not walk between one walking bout and the next, averaged for all pauses.

**Statistical analysis**

All statistical tests were conducted with MATLAB. To compare the order parameters, Wilcoxon rank-sum test was used. To compare among the three stages, Friedman Test was used. Since all comparisons were significant according to the latter test, Bonferroni multiple comparison post-hoc test accompanied each Friedman Test, in order to reveal the significantly different groups. The $p$ values noted are of the Bonferroni multiple comparison post-hoc test. Correlations were conducted using Spearman's Rank Correlation test.

**Modelling**

We employed a two-dimensional model comprising $N$ agents moving at a fixed speed $v$ in a square domain of linear size $L$ and periodic boundaries. Position and heading (direction) of each rectangle centre were denoted by $x_i(t)$ and $\hat{v}_i(t)$, respectively, where

$x_i(t) \in [0, L]^2$ and $\|\hat{v}_i(t)\| = 1$. In addition, $w_i(t)$ is a Boolean variable that denoted whether at time *t*, agent *i* is moving ($w_i(t) = 1$) or pausing ($w_i(t) = 0$). In order to take into account the physical size of animals, we assumed that each agent is a rectangle with sides $a_1 \times a_2$.

In each simulation step, the position of moving agents changed by amount *v* in the direction $\hat{v}_i(t)$,

$$x_i(t+1) = x_i(t) + v w_i(t) \hat{v}_i(t) \quad (mod\ L).$$

The new velocity of agent $i$ depended on three terms: i. Inertia (its own direction); ii. the direction of other agents in its receptive field (see below), possibly with a memory; and iii. a random vector with unit norm $\xi_i(t)$. The receptive field view is a circle around $x_i(t)$. Unlike previous models, we assumed that an agent sees all other agents whose rectangle is within a distance *r* from $x_i(t)$, and not merely the center of the rectangle. Denoting the four corners of the rectangle describing agent *j* as indexes $c_j^1(t), \ldots, c_j^4(t)$, we defined the edges of the rectangle as the line segment $e_j^k(t) = [c_j^k(t), c_j^{k+1}(t)]$, where $c_j^5(t) = c_j^1(t)$. The adjacency matrix $A_{ij}(t)$ is a zero-one matrix that is one if agent *j* can be observed by agent *i* at time *t*, i.e.,

$$A_{ij}(t) = \begin{cases} 1 & \left(e_j^1(t) \cup e_j^2(t) \cup e_j^3(t) \cup e_j^4(t)\right) \cap \{x: \|x_i(t) - x\| \le r\} \ne \phi \\ 0 & \text{otherwise} \end{cases}$$

Then, the set of observable neighbours of agent *i* at time *t* as,

$$I_i(t) \coloneqq \sum_{i \ne j} A_{ij}(t).$$

The average direction in the receptive field at time $t$, is $\hat{u}_i(t) = u_i(t)/\|u_i(t)\|$, where

$$u_i(t) = \left(\frac{1}{n_i(t)}\sum_{j\in I_i(t)} \cos(\hat{v}_j), \frac{1}{n_i(t)}\sum_{j\in I_i(t)} \sin(\hat{v}_j)\right).$$

Here, $n_i(t)$ is the number of elements in $I_i(t)$.

Overall, the new heading $\hat{v}_i(t+1)$ was the weighted average of four terms: inertia, direction of moving intersecting rectangles at time $t$, the average heading of moving intersecting rectangles at time $[t-5 \ldots t-1]$, and independent random vector with unit norm $\xi_i(t)$ from a uniform distribution of $[-\pi, \pi]$. Angles of other agents to the focal agent were provided by the simulator. Let the set of the weights be $[q_1 \ldots q_4]$.The updated weighted averaged direction, was, therefore, $\hat{v}_i(t) = v_i(t)/\|v_i(t)\|$, where

$$v_i(t+1) = q_1\hat{v}_i(t) + q_2\hat{u}_i(t) + q_3\frac{1}{5}\sum_{s=1}^{5}\hat{u}(t-s) + q_4\xi_i(t)$$

The weights were set to $Q = [0.2, 0.2, 0.3, 0.3]$. Collisions of agents with each other were not taken into account, as our previous study found that visual stimuli is sufficient for generating collective behaviour (Ariel et al., 2014a).

Switching between walking and pausing was determined as follows. Assuming exponential distributions of moving and pausing durations, the probability to proceed moving was given by a single parameter, $p_w$, describing the probability per-step of an agent to continue walking, $p_w = P(w_i(t+1) = 1 | w_i(t) = 1)$. Similarly, $p_s = P(w_i(t+1) = 0 | w_i(t) = 0)$ was the probability per-step that a stopping agent will continue stopping.

The following statistics were calculated over steps 200-1000 of the simulation. Brackets denote averaging over all analysed frames.

1. Order: $\theta(t) = \frac{1}{N}\langle\left\|\sum_{i=1}^{N}\hat{v}_i\right\|\rangle$ .

2. Spread: $S(t) = \frac{1}{N(N-1)} \langle \sum_{i=1}^{N} \sum_{j \neq i} \left| x_i(t) - x_j(t) \right| \rangle$
3. Number of neighbours: $n_i(t) = \frac{1}{N} \langle \sum_{i=1}^{N} I_i(t) \rangle$
4. Steps to regroup: the number of steps that passed from the first step in which an agent has no neighbours within its visual field $(n_i(t) = 0)$ until it encounters a neighbour again $(n_i(t+s) > 0)$.
